# ATI_Box: A Simple tool for convolutional neural network-based image semantic segmentation

**DOI:** 10.64898/2026.05.29.728143

**Authors:** Tomasz Przygodzki

**Affiliations:** Department of Haemostatic Disorders, Chair of Biomedical Sciences, Medical University of Lodz, ul. Mazowiecka 6/8, Lodz, 92-215, Poland

## Abstract

Quantitative analysis of microscopic images has become a standard in basic biological and biomedical research. Deep machine learning provided a powerful tool facilitating this process. However, practical adoption of deep machine learning to image analysis may be difficult for a researcher who lacks basic coding skills. This is caused by a limited number of non-coding solutions, specifically in the domain of convolutional neural networks (CNNs). This scarcity may be explained by the following paradox. Training of CNNs is a relatively complex process. Researchers who are familiar with this process are also skilled enough to code the full pipeline of CNN implementation from annotation, through model training and evaluation to its usage in laboratory practice. Any kind of an alternative solution, acceptable by a broader group of researchers who are unfamiliar with CNN concepts, must inevitably result in simplification of the entire process, specifically the training step. Such simplification in turn may lead to limitation to solve specific problems by such a tool. Author believes however, that some compromise may be found between complexity and simplicity that would be sufficient to solve some basic problems in the field of basic biological and biomedical research.

To address this challenge, author proposes ATI_Box (Annotation, Training, Inference in One Box), a unified, user-oriented platform for end-to-end image semantic segmentation. The system integrates data annotation, storage, model training, evaluation, and quantitative analysis into a single workflow, significantly simplifying the model development process. Image and annotation data are managed through an S3-compatible object storage system (MinIO), enabling scalable and transparent data handling. Annotation process is implemented through Label Studio. Model training is based on convolutional neural network U-Net architecture with ResNet as an encoder. Model evaluation is performed on ground-truth dataset held-out during training and provides pixel-level and object-level evaluation metrics. Batch analysis mode enables automated quantification of model predictions such as object counts and coverage areas. The usability of the platform was presented on examples from laboratory practice.

The platform is intentionally devoid of model-tuning capabilities as it is addressed to users unfamiliar with profound machine learning concepts. At the same time, accessibility of such basic features of model training as definition of epochs number or saving and implementing of trained model versions enables one to perform some basic analytical experiments. As such, the platform may serve not only as an analytical tool but also as an educational solution to explain practical basics of semantic segmentation process.

## 1. Introduction

### 1.1 Background

Microscopy is one of the leading methods in the field of basic biological and biomedical research. Its role is no longer limited to visualisation and presentation of qualitative features, but it became a powerful tool for quantitative analysis. Specific tasks include, but are not limited to counting cells, measuring their area, distinguishing cells of various morphological classes, or quantifying subcellular structures.

For a long time, automated analysis of digital images were based on relatively simple mathematical operations on pixel intensities. Nowadays, with the advancement of machine learning methods, the analysis became more complex and effective. One of the implementations of machine learning in image analysis is discerning objects which belong to different classes – a process called semantic segmentation. It allows automatic quantification of structures with respect to their type defined by an analyst. There are several machine learning methods which allow semantic segmentation: random forest classifier, convolutional neural networks and vision transformers. The first two methods are most often used for semantic segmentation tasks in analysis of microscopic images.

### 1.2 Motivation for implementation of an integrated platform for semantic segmentation

The use of machine learning methods is a process that leads through training of a model on a set of data, evaluation of the trained model and finally its usage in routine work. Training and application of convolutional neural networks for semantic segmentation, like many other procedures which involve deep machine learning, is a multistep process. Consecutive steps are implemented by scripts, typically written in Python. The necessity of writing and executing a code creates too high barrier to entry for many researchers. Therefore, some no-coding solutions which cover the whole pipeline of semantic segmentation from image annotation to quantitative data are being developed. The tools that, to the best of author’s knowledge, come closest to this concept are briefly described below and listed in **Table 1**, along with an indication of the extent to which they perform the selected functions.

**Table 1.** Brief summary of selected tools for machine learning-based image analysis.

| Tool name | Annotation | CNN training<br>on user-<br>annotated<br>data | Model<br>evaluation | Batch<br>quantification | Single<br>GUI | No-code<br>solution |
| --- | --- | --- | --- | --- | --- | --- |
| Ilastik[1] | + | -<br>(random forest<br>Classifier;<br>inference with<br>pretrained CNN<br>models) | + | + | + | + |
| QuPath (native)[2] | + | -<br>(random forest /<br>scripting for<br>CNN) | + | + | + | + |
| CellProfiler<br>(+analyst)[3, 4] | -<br>(automatic<br>object<br>detection) | - | + | + | + | + |
| ZeroCostDL4Mic[5] | - | + | + | + | -<br>(Jupyter<br>notebooks) | + |
| DeepMIB[6] | + | + | + | + | + | + |
| ATI_BOX | + | + | + | + | + | + |

Ilastik is a package for segmentation based on random forest classification algorithm. It has intuitive GUI and allows to perform analysis from uploading images, through classifier training to quantitative results. It has all advantages of random forest classifier i.e. a fast training (effects noticed in real time) and no requirements for GPU. It may be less effective when complex structures defined by long-range features are to be segmented. At the same time it allows inference with the use of pretrained CNN models. QuPath is a package that allows cell annotation, segmentation and classification based on intensity thresholding and watershed algorithm combined with random forest classifier. It is designed for detecting cells and nuclei. It was created for whole slide analysis and is very often used for this purpose. CellProfiler Analyst allows training of classifier based on crops of images containing objects of interest. Classification uses a plethora of machine learning methods. Since it does not have a manual annotation tool to delineate object boundaries it is designed to detect separate object rather than complex, interweaving structures. ZeroCostDL4Mic is a framework which allows defined operations through Jupyter notebooks on Google Colab. These operations include, among others, objects segmentation with the use of U-Net. The platform does not provide its own image annotation tool. DeepMIB is the solution which fulfils all the criteria of annotation, training of CNN models, their evaluation and implementation in one place. Notably, it provides robust capabilities of the choice of CNN architecture and model tuning. Its broad capabilities, on the other hand, may be overwhelming for entry users.

In this paper yet another solution for CNN-based semantic segmentation is proposed - ATI_Box (Annotation, Training, Inference in One Box). The platform was created as an in-house tool for semantic segmentation, covering the entire process - from image annotation, through model training and evaluation, to its implementation in routine analysis - in the simplest possible way. The advantage of such simplified tool, and at the same time the author’s motivation for creating it, was to enable a larger group of researchers to assess the usefulness of CNN-based solutions for their routine analysis. Although this simplification comes at the cost of limiting the full capabilities of CNNs, the author believes that such a solution would still be sufficient to solve basic problems in the field of basic biological research.

## 2. Methodology

### 2.1 Code generation

A notable dimension of ATI_Box is that its implementation was largely enabled by generative AI– assisted programming tool Claude (Anthropic, model version: claude-sonnet-4-6). As a trained biologist rather than computer scientist, the author possesses basic Python skills but lacks formal software engineering expertise. Therefore the author took the advantage of generative AI as an implementation partner, translating the architectural intentions into executable code through iterative refinement and validation.

### 2.2 Platform architecture

#### 2.2.1 General description

ATI_Box has modular architecture. The main user interface is implemented using the Streamlit framework, which provides web-based environment through which users interact with all stages of the segmentation pipeline, including image upload, project configuration, model training and evaluation and inference.

The platform is implemented as a multi-container system orchestrated using Docker. It consists of three main services: (i) a core application container hosting the user interface and machine learning pipelines, (ii) a Label Studio container providing annotation capabilities, and (iii) a MinIO container serving as an S3-compatible object storage backend. These services communicate using application programming interfaces (**Figure 1**).

**Figure 1.**
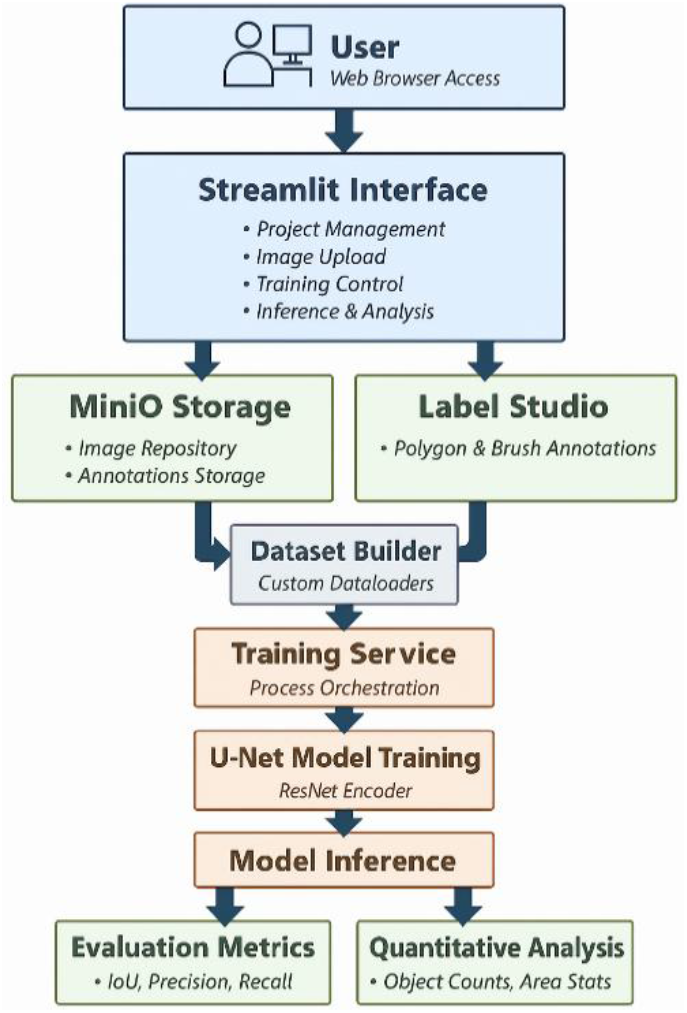
Schematic representation of ATI_Box architecture.

#### 2.2.2 Data Storage and Management

Images can be uploaded in any of the graphical formats such as tif, jpg or png with the latter being the format used effectively in the pipeline. Images in other formats are automatically converted to png. Uploaded images are stored in the MinIO object storage system. The storage layer serves as the central data repository of the platform, providing access to datasets used during annotation, training, and model evaluation. The use of an S3-compatible object storage system reflects a standard architectural requirement shared by contemporary annotation platforms, including Label Studio which is used as annotation tool in ATI_Box. Folders used by the platform (Label Studio data, MinIO storage and model checkpoints) are located on the host machine and mounted into containers using bind mounts. This design enables direct access to data through the host filesystem. This facilitates data portability, allowing datasets, annotations, and trained models to be easily transferred between machines without requiring Docker-specific export procedures. Although bind mounts require a predefined directory structure on the host system, this is automatically created by initialization scripts (start.sh and start.bat) itself, ensuring consistency across deployments.

#### 2.2.3 Annotation Workflow

Image annotation is performed using Label Studio, which is integrated into the platform. Users define the number of segmentation classes as well their names and choose the annotation mode (polygon or brush) in Streamlit interface (**Figure 2**). This results in generation and configuration of Label Studio project through its API. The system opens the annotation interface directly in Label Studio. Images stored in MinIO are automatically registered as annotation tasks within the project, allowing them to appear immediately in the annotation interface. When annotation is completed, the platform retrieves the generated labels from Label Studio and converts them into segmentation masks that can be used for model training.

**Figure 2.**
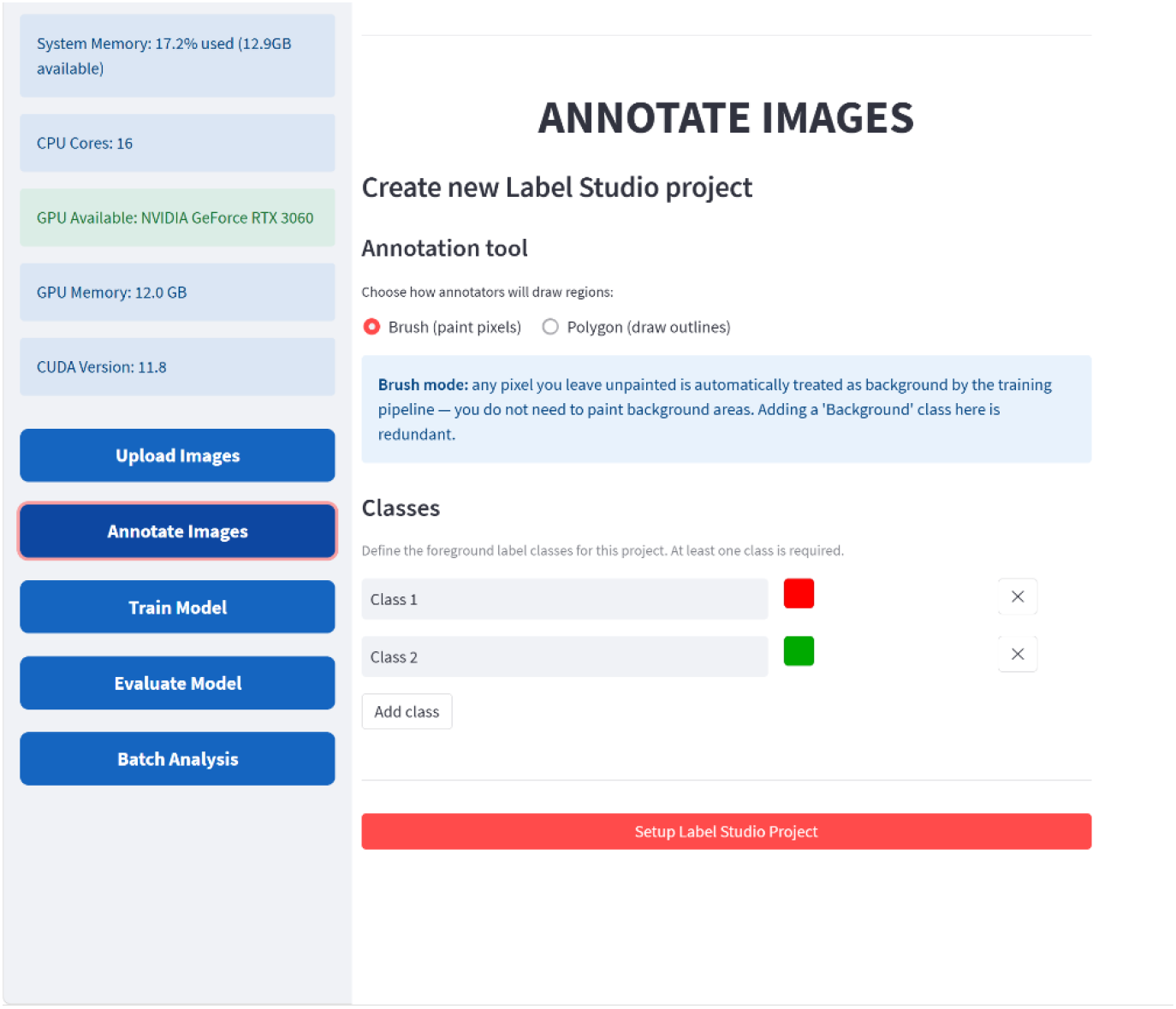
Graphical user interface which allows configuration of Label Studio. User chooses annotation type, brush or polygon mode, and defines number and names of classes to be annotated.

#### 2.2.4 Model Training

Model training is performed using annotated images retrieved from the storage layer and converted into segmentation masks. The training procedure is implemented in PyTorch and executed as a separate background process in order to prevent blocking the user interface. During initialization, the system determines the training configuration based on detected device characteristics. Hardware properties, including GPU availability, device name, total memory, and CUDA version, are identified using PyTorch (via torch.cuda). These parameters are subsequently used to adjust selected training settings through predefined rules: batch size is scaled according to available GPU memory, while encoder selection follows a fixed strategy based on device type, with ResNet101 used when a GPU is available and ResNet50 otherwise. Additionally, device-specific configurations, including gradient checkpointing for GPU execution and environment variable adjustments, are applied to ensure efficient resource utilization.

The segmentation model is based on the U-Net architecture implemented using the segmentation_models_pytorch library [7]. U-Net was selected because it has become one of the most widely used architectures for biomedical image segmentation due to its ability to combine high-resolution spatial information from early network layers with high-level semantic features extracted by deeper layers. This architecture has proven particularly effective for microscopy images where precise localization of object boundaries is required [8-10]. The encoder backbone is selected automatically according to the available hardware resources, while the number of output channels is determined dynamically from the annotation classes detected in the project configuration.

Images from the training set and their corresponding annotation masks are loaded directly from the object storage system. Custom dataset loaders are used to retrieve images and annotations generated by the annotation interface and convert them into numerical mask representations. Prior to training, a fixed held-out test set is created by randomly selecting 10% of annotated images (with a minimum of one image), which are persistently stored and excluded from all subsequent training and validation procedures. These images and masks are to be used as ground truth for model evaluation after completion of training. The remaining data are then divided into training and validation subsets using an 80:20 split. Input images are resized to 512 × 512 pixels and augmented using geometric transformations including horizontal and vertical flipping, and random 90-degree rotations. These augmentations increase the effective diversity of the training dataset and improve model generalization. Images are subsequently normalized and converted into tensor representations before being passed to the neural network. Mini-batches are generated using PyTorch DataLoader objects. The batch size is determined automatically based on the detected compute device and available memory resources. When GPU acceleration is available, multiple worker processes and pinned memory are used to accelerate data loading, whereas CPU-based training uses a simplified configuration to reduce memory consumption.

Model optimization is performed using the Adam optimizer with a learning rate of 1 × 10^−4^. The training objective is defined as a combined loss function consisting of focal loss and Dice loss. The combination of two losses has been proposed as a way to improve segmentation performance in cases where objects occupy only a small fraction of the image area, which is common in biological microscopy datasets [11].

User defines the number of training epochs and default number is set at 100. During each epoch the model processes all batches in the training dataset, performing forward propagation, loss computation, gradient backpropagation, and parameter updates. Intermediate tensors are explicitly dereferenced after each training batch, and memory cleanup procedures, including Python garbage collection and GPU cache clearing, are invoked to reduce memory consumption. This approach mitigates memory accumulation and fragmentation, which is particularly important in containerized environments with limited resources.

After completing the training phase of each epoch the model is evaluated on the validation dataset. Validation is performed with gradient computation disabled and computes the average validation loss across all validation batches. Both training and validation losses are reported after each epoch, allowing monitoring of model convergence and early identification of potential overfitting. Training and validation loss curves are shown as training progresses (**Figure 3**.).

**Figure 3.**
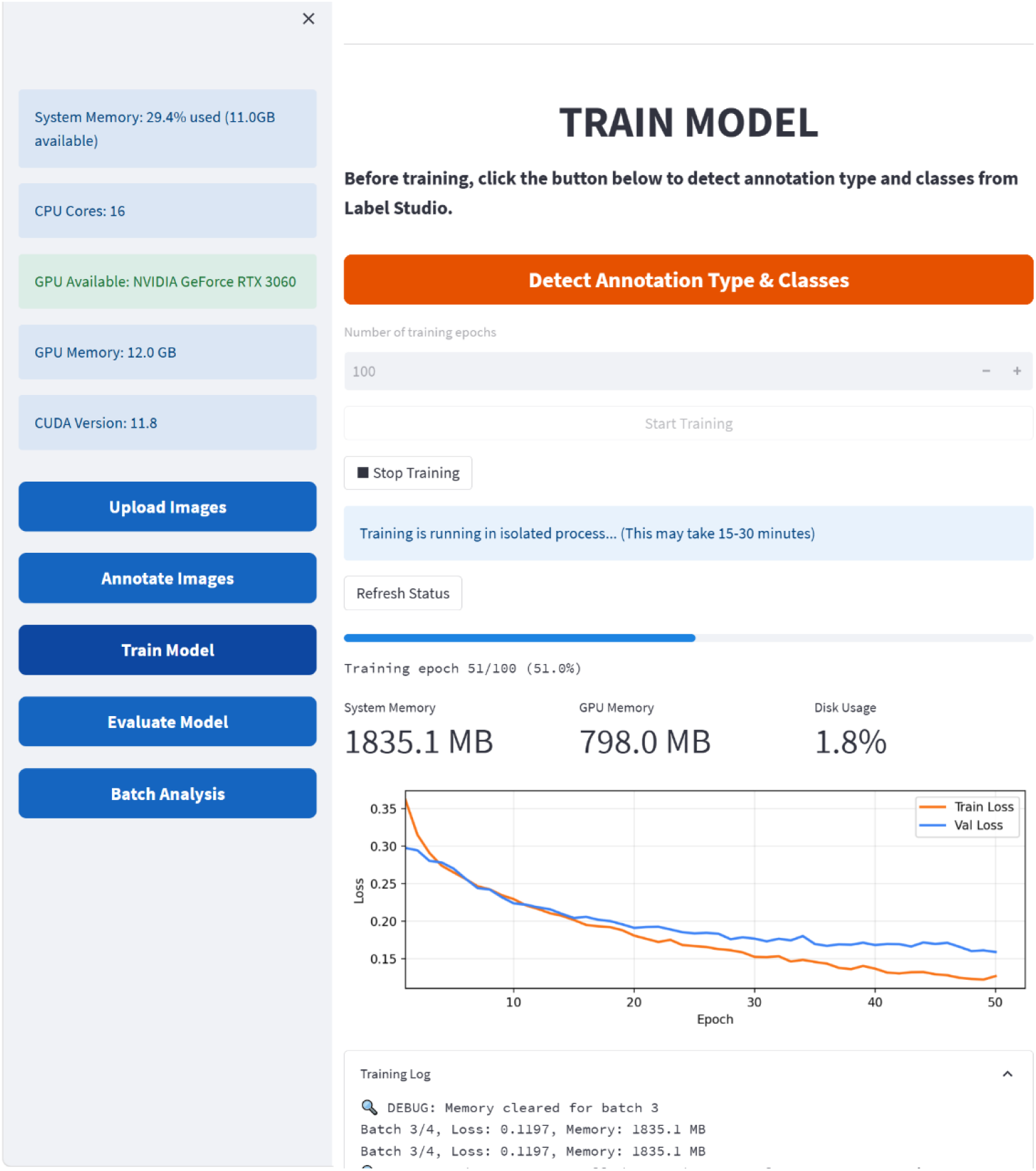
Graphical user interface for model training. User defines number of epochs. Progress of training is displayed along with train and validation loss curves.

To reduce disk usage within the containerized training environment, intermediate checkpoints are not stored during training. Instead, only the final trained model is saved upon completion of the training process. The saved model is accompanied by a configuration file that records the encoder architecture, annotation mode, and class definitions, ensuring that the model can be correctly reloaded for subsequent inference and batch analysis tasks.

#### 2.2.5 Inference and Quantitative Analysis

The platform provides several modes for applying the trained model and analysing segmentation results. These include model evaluation on a single image, evaluation against annotated test images stored in Label Studio (ground truth), and finally the actual usage i.e. batch quantitative measurements on research data.

Implementation of the inference modules is shown with the use of a real example. The model was trained to differentiate round objects (procoagulant platelets) from a homogenous mass of platelets forming thrombi in microfluidic chamber system. This task will be described in details in **Results** section.

##### Single-Image Inference

Inference on a single image is intended as a quick visual inspection of the model convergence. The user selects a version of a trained model and uploads an image, which is automatically converted to PNG format if necessary. The image is processed by the trained segmentation model, producing masks for each predicted class (**Figure 4**.).

**Figure 4.**
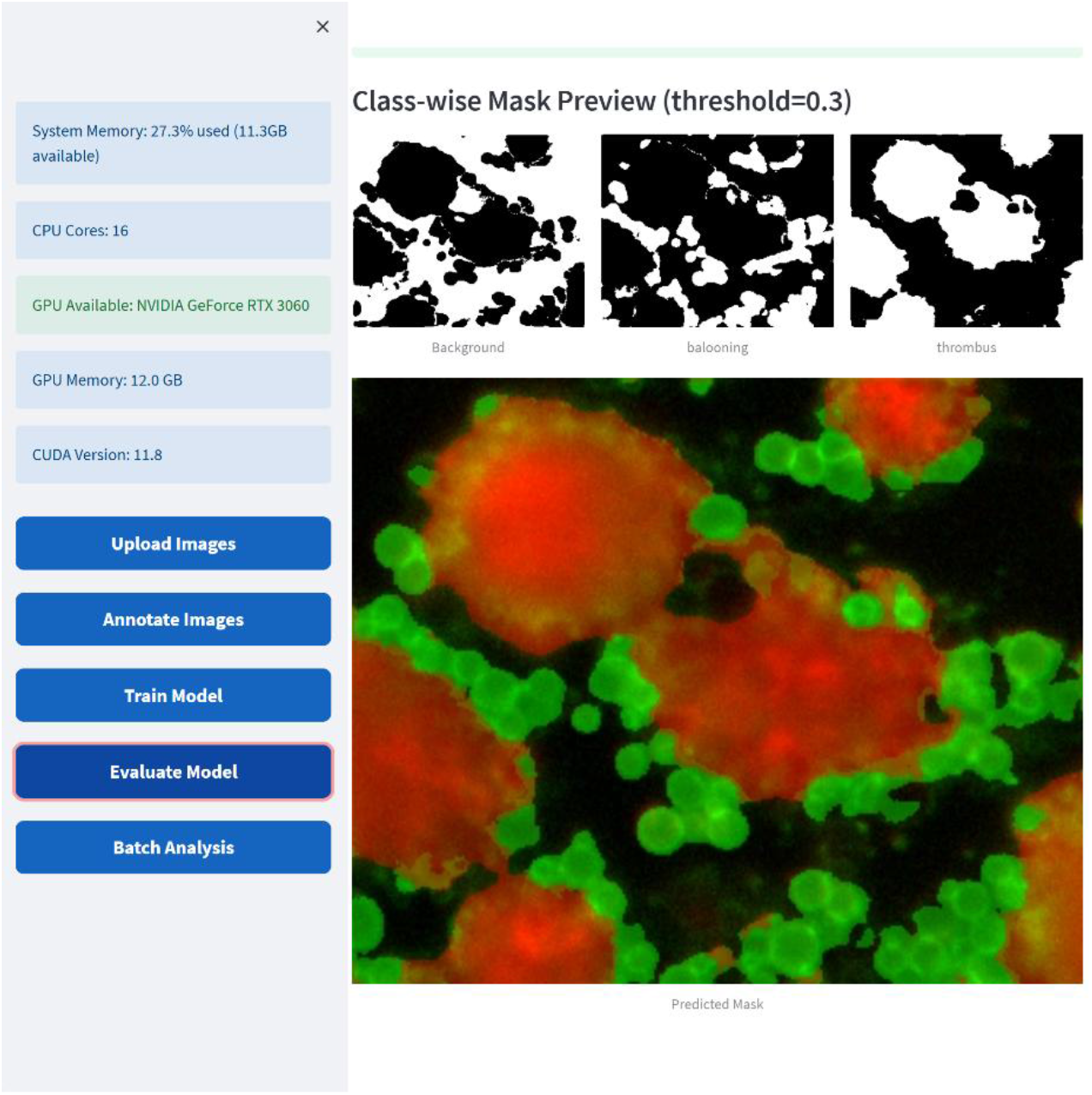
Inference on single image. For a quick visual inspection of the model convergence an inference on a newly uploaded image can be performed with a probability threshold adjusted to a desired level.

The application visualizes the results by generating an overlay in which predicted segmentation masks are displayed on top of the original image using class-specific colours. Individual class masks are also displayed separately to allow inspection of the model’s predictions for each category. The segmentation threshold used to convert model probabilities into binary masks can be adjusted interactively through a user-controlled parameter, allowing users to explore how the prediction confidence influences segmentation results. This mode is intended primarily for rapid qualitative assessment of model performance and for visual inspection of predictions on new images.

##### Evaluation Using Ground Truth from Label Studio

Prior to model training, the system automatically creates a held-out test set by randomly selecting a subset of annotated images (10%, with a minimum of one image) from the dataset as described in Model Training section. These images are excluded from all subsequent training and validation procedures and are persistently stored to ensure consistency across runs. During evaluation, this fixed test subset is used for batch inference, providing an unbiased assessment of model performance on data never seen during training. The system retrieves annotated images and their corresponding segmentation masks from the storage backend and performs automated batch evaluation.

For each image, the model generates predicted segmentation masks that are compared with the ground-truth masks derived from the annotations. The evaluation calculates several quantitative performance metrics, including pixel-level intersection over union (IOU) for each segmentation class, mean IOU averaged across classes, object-wise precision, object-wise recall and object-wise F1-score.

Object-wise metrics are computed by identifying connected components in both predicted and ground-truth masks and matching predicted objects to annotated objects using a defined IoU threshold value of 0.5. This evaluation allows assessment not only of pixel accuracy but also of the model’s ability to correctly detect individual objects. The results are presented in the user interface as summary metrics together with detailed per-class performance scores (**Figure 5**).

**Figure 5.**
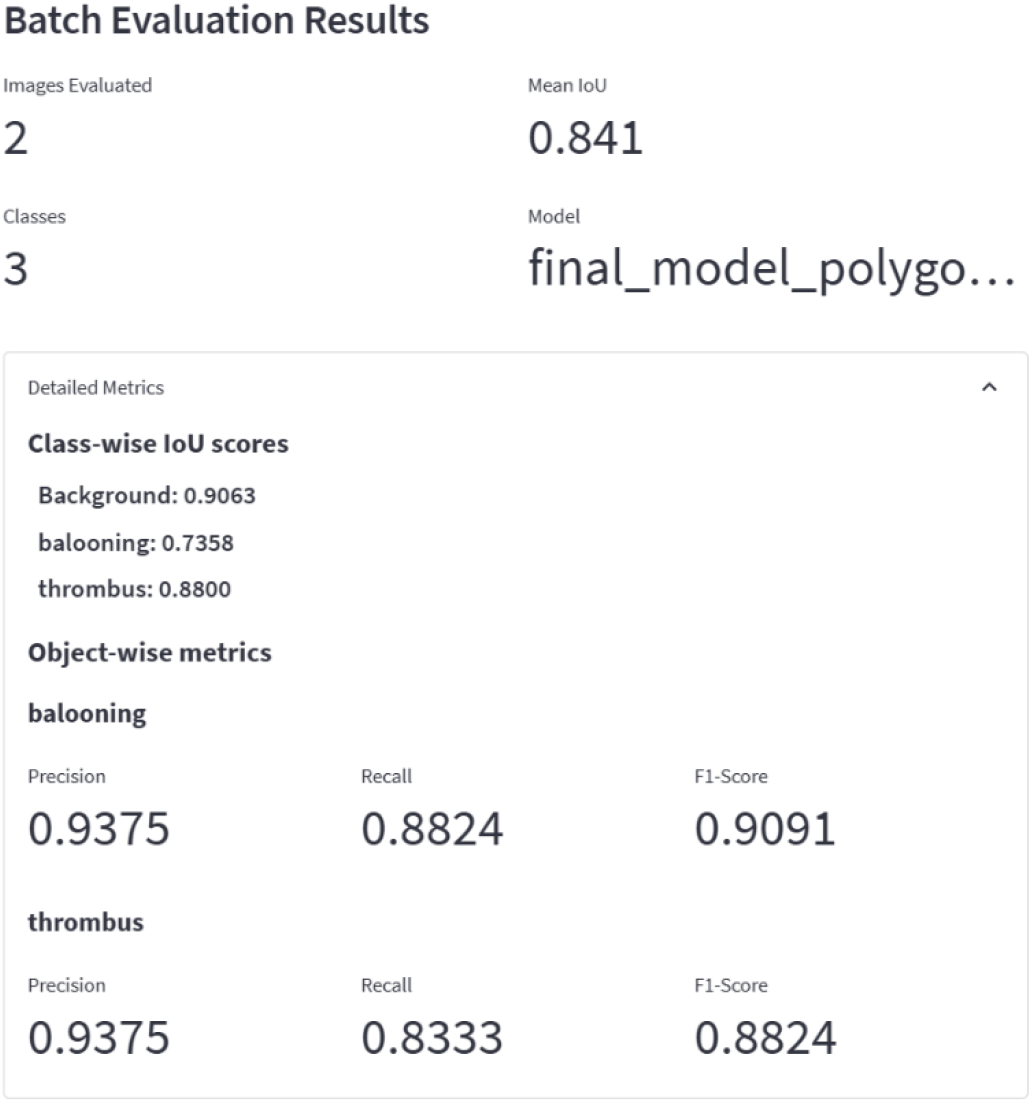
Inference on ground truth images. Model inference is performed on ground truth images not used in training process. Pixel-wise and object-wise metrics are calculated.

##### Batch Analysis and Quantitative Measurements

When model evaluation provides satisfactory convergence, users can apply it for quantitative evaluation of their actual experimental data. The platform includes a batch analysis module designed for extracting quantitative measurements from segmentation results. In this mode a user uploads a set of images that are converted to png format if needed and processed automatically using the trained model. Besides selecting segmentation threshold of probability, a user can define a size threshold to filter out objects smaller than defined value if, based on user’s expertise, they are irrelevant (**Figure 6**).

**Figure 6.**
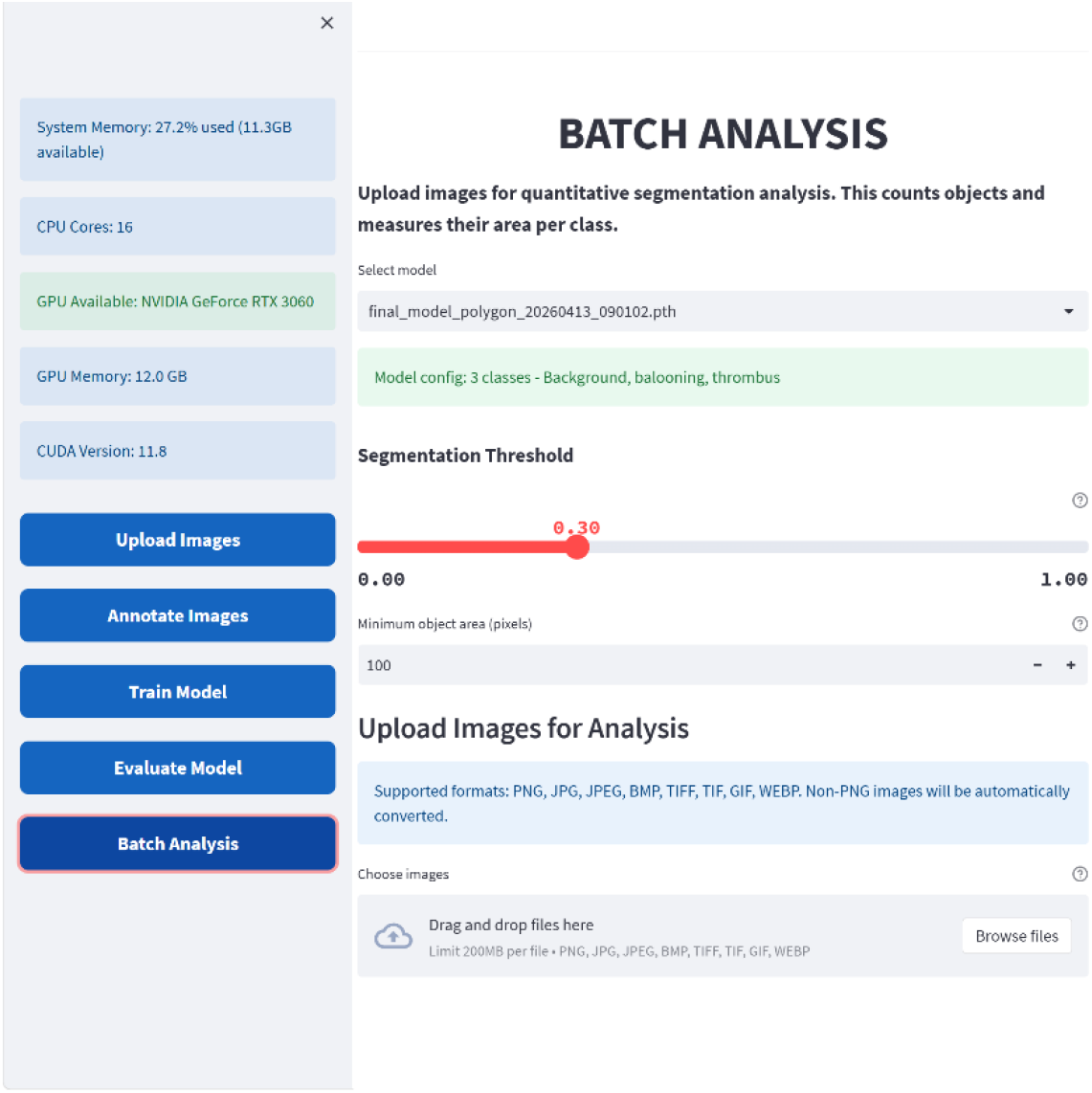
Inference on a batch of images. Model inference is performed on batch of uploaded images with defined probability and size threshold.

The analysis computes several metrics for each class, including: number of detected objects, total segmented area in pixels and percentage of image area occupied by the class.

The results are shown in a table for all images in the set (**Figure 7**).

**Figure 7.**
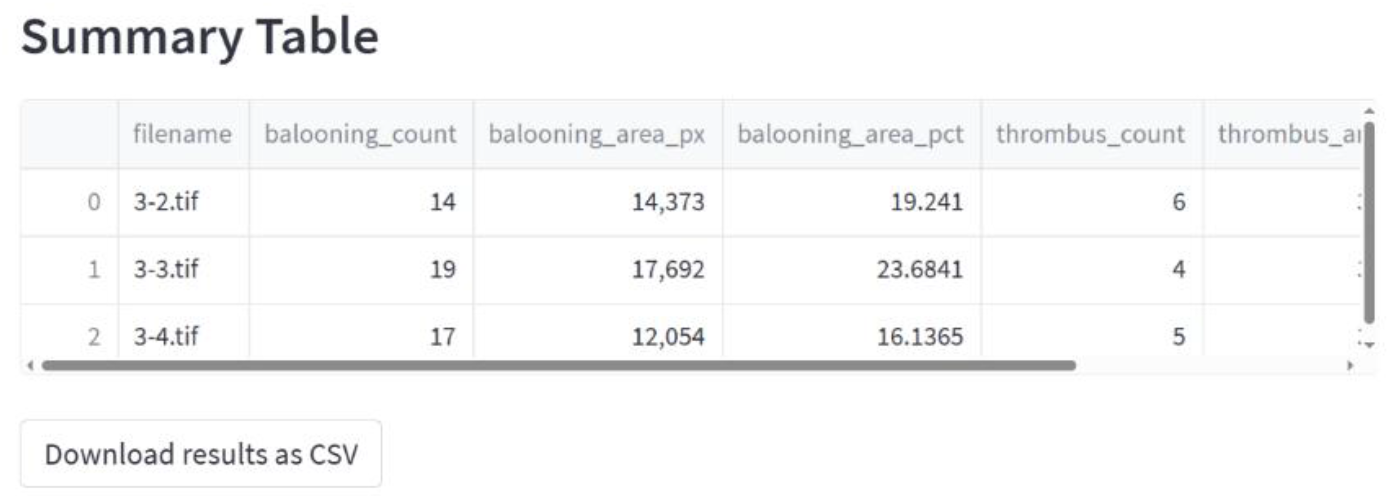
Combined results display. Results of analysis are displayed for the batch of images. For each image in the batch number and area of objects of each class are displayed in a table.

The application also generates visual overlays showing the predicted segmentation masks for each image in the set. These overlays allow users to visually verify the segmentation results and the detected objects (**Figure 8**).

**Figure 8.**
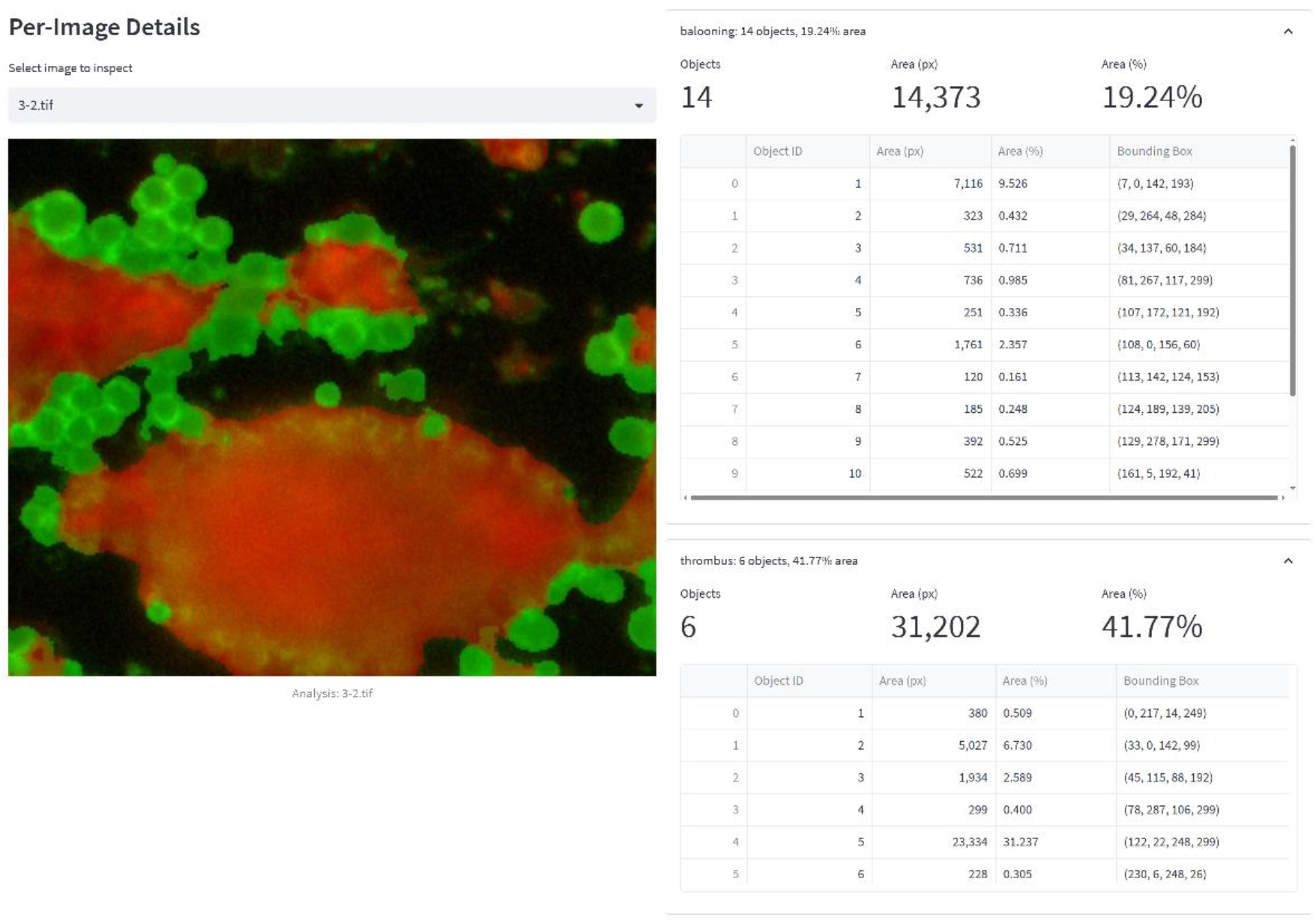
Inference results for a single image from a batch. Results and overlying masks can be inspected for each image in the batch separately.

The platform aggregates the results into a summary table containing one row per image and columns describing object counts and area measurements for each class. The results can be exported as a CSV file, enabling downstream statistical analysis in external tools such as Python, R, or spreadsheet software.

#### 2.2.6 Projects separation

The platform supports persistent storage of multiple datasets and annotation projects; however, the user interface is designed to operate on a single active project at a time. Once a project is initialized, its identifier is stored in a configuration file and automatically reloaded on subsequent sessions, effectively binding the application to that project. To create a new project a user needs to create folder for this project and recreate in this folder the installation procedure to deploy a new container. This will result in label-studio-data and minio-data folders creation where images and annotations for this project will be stored. It is assumed that user would work only on one project at a time on a machine, therefore all containers share the same set of ports. The most convenient way for the user to switch between projects is to stop the container with a project that users ceases to work and to start the one that user chooses to work with. This can be done either in Docker Desktop or by command line by “compose down” in the currently used project and “compose up” in the folder where the new project resides.

## 3. Results

To show ATI_Box usability, three exemplary implementations are shown that may be considered representative for a variety of tasks met in the laboratory practice.

### 3.1 Quantification of procoagulant platelets in thrombi formed in flow chamber assay – example from the fluorescence microscopy field

Data from this experiment were shown in the previous section to illustrate ATI_Box interface. Blood platelets which form thrombi on collagen in flow chamber assay present two morphological types: one of them is a merely homogenous mass of aggregated platelets and the other are platelets which undergo a transition to procoagulant form characterized, among other features, by “ballooned” shape. The task was to quantify these structures. The following criterion was established to classify objects: only platelets with recognizable entire circumference were annotated as procoagulant platelets, remaining structures were annotated as thrombi (**Figure 9 a,b**). The dataset consisted of twenty images. Model was trained for 100 epochs. Evaluation metrics on ground truth data (2 images held-out from training dataset) are presented in **Figure 5** in the above section. Although the held-out set of 2 images is too small for robust performance estimation it provides preliminary evidence of model convergence. The model was used to quantify differences in the area covered by procoagulant platelets in the time course of thrombus formation (**Figure 9 c-f**).

**Figure 9.**
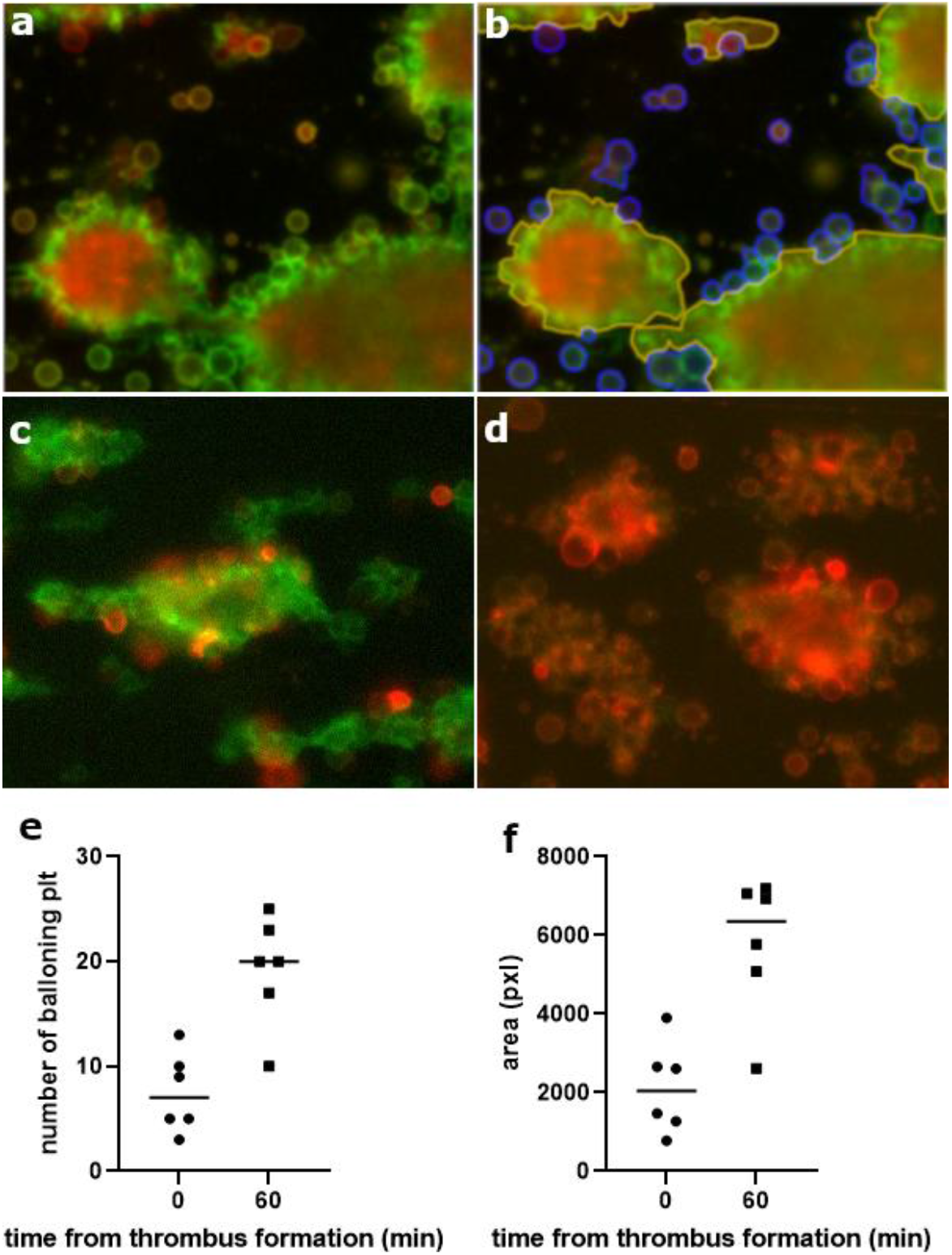
Quantification of ballooning blood platelets. (a) Exemplary image of thrombus formed on collagen and its annotations (b), yellow lines used for thrombi, blue lines - procoagulant platelets. Images of thrombi shortly after formation (c), and 60 minutes after formation (d), presenting low and high number of procoagulant platelets respectively. Results of quantification of six images with trained model – (e) number of identified ballooning platelets and (f) area covered by this class, each dot represents results calculated for one image

### 3.2 Quantification of wound in scratch assay – “cell culture room classics”

A scratch assay or wound healing assay is a very often used assessment of cell migration. In a dish with cell monolayer a scratch is made which mechanically removes portion of cells and the ability of remaining cells to cover the gap by proliferation and migration is assessed. To this end an area or the width of the “wound” is measured. This is done either manually or with the assistance of dedicated image analysis tools. Two most often used solutions are MiToBo Scratch Assay Analyzer [12] and *Wound_healing_size_tool* plugin for ImageJ [13] which can be both used as ImageJ plugins. These tools are reported effective by many users. In problematic cases such as low contrast images or uneven illumination CNN-based solution may be of some help. ATI_Box was tested on series of images characterized by low contrast and uneven edges of a wound (**Figure 10**). Model was trained to segment area covered by cells rather than a wound itself as the latter is a plain plastic surface devoid of patterns on which the model can efficiently converge. Since the analysis module calculates the percentage of the area covered by each class, an increase in area covered by cells may be used as a measure of wound closure. As shown on Figure 10 the model was trained on 10 images.

**Figure 10.**
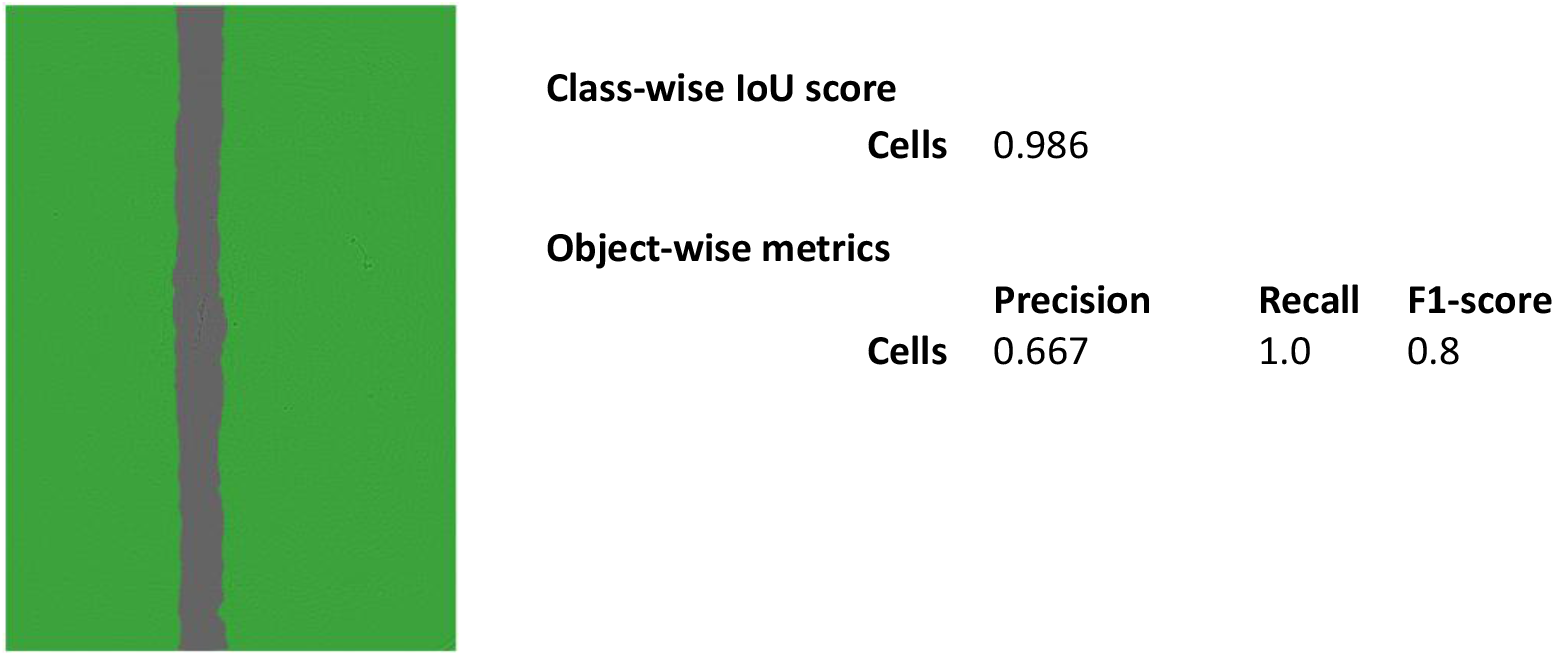
Segmentation of cell monolayer surrounding artificial wound in scratch assay. Left panel presents cell monolayer segmented by the trained model (green mask). Model evaluation metrics calculated for 2 images from a test set.

Relatively low precision is plausibly due to the fact, that images which were selected by training script for model evaluation contained cell-free patches located outside the wound and annotated as a monolayer. Since these patches are not a part of the wound their area obviously should not be added to the wound area. If they are detected by the model they can be filtered out with size threshold in batch analysis mode.

### 3.3 Quantification of blood morphological elements – example from bright-field microscopy

Recognition of red and white blood cells as well as blood platelets in blood smears was used to show an example of ATI_Box performance on bright-field images. It was also used to test the way to overcome of a problem of annotation of images with large number of objects with highly imbalanced classes. As shown on **Figure 11a** the smears present large number of red blood cells (RBCs), much smaller number of platelets and single instances of leukocytes. Annotation of all RBC instances present on image is extremely laborious task and also introduces high class imbalance. At the same time annotation of only selected instances of the class would result in an ineffective training. One possible way to overcome this problem is to train a model on image crops which contain less objects and where the classes are better balanced. This approach turned out to be ineffective. Although the model trained on crops had good convergence when tested on cropped images (**Figure 11 b,c**), its inference on full size image was extremely poor (**Figure 11d**). Therefore another approach was used where full-sized images were patched with artificial background crops. In this way majority of objects was obscured and only area with low number of objects of each class were left which resulted in better balanced representation of each class (**Figure 11e**). Model training on such prepared and annotated images led to much better model convergence on full size images of smear (**Figure 11f**).

**Figure 11.**
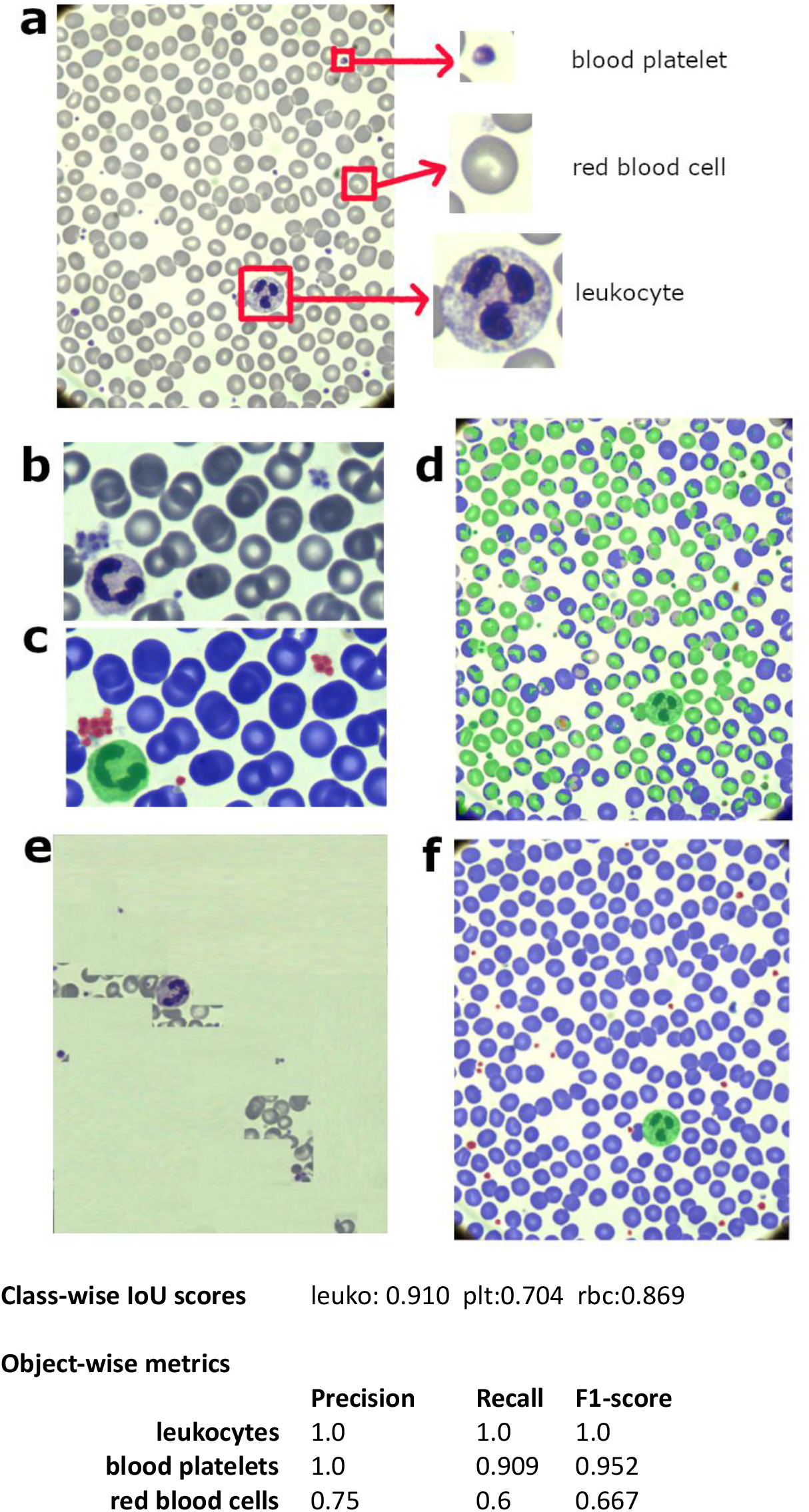
Segmentation of morphological blood elements. (a) exemplary blood smear, (b) cropped smear image, (c) inference of a cropped image with the use of a model trained on cropped fragments, (d) inference of a full-size image with the use of a model trained on cropped fragments, (e) full size image patched with artificial background fragments, (f) inference of a full-size image with the use of a model trained on full size image patched with artificial background fragments. Inference masks colouring scheme: green-leukocytes, red – blood platelets, blue-red blood cells

What draws attention in evaluation metrics is relatively low recall and precision for red blood cells class despite very good segmentation of this class as evidenced by both inference image and by class-wise IoU score. This is due to the way the object-wise metrics are calculated in case of touching objects. Many red blood cells touch each other. Some of them could have been annotated as separated objects, but detected by the model as one merged object. In such case only one annotated object was calculated as true positive, while the other one was treated as false negative (it was not detected by the model as a separate object). Therefore, if the number of red blood cells was the crucial value in the experiment, the researcher would have to increase number of images in the training set or to improve annotation of red blood cells and retrain the model.

## 4. Discussion

Although the programming skills and concepts of machine learning increase among researchers working in biological or biomedical field, the necessity of writing and executing a code still creates too high barrier to entry for many of them. Some of the researchers may benefit from an access to facilities specializing in acquisition and analysis of microscopic images in their institutions, but it applies only to selected universities. Increasing number of free of charge, open code solutions is progressively filling this gap. As described above, some of them are friendly even for entry users while others are robust solutions which may be overwhelming for a novice. In this context, ATI_box is positioned as a tool which can help users who have not yet had the opportunity to work with CNN-based image segmentation and analysis become familiar with this kind of analysis.

Undoubtedly ATI_Box oversimplifies CNN training process by depriving it from hyperparameter tuning. At the same time though it is capable of solving basic segmentation tasks as shown by exemplary implementations. An experience of using such a basic tool can also stimulate some of its users to dive deeper into the machine learning concepts. It is facilitated by the fact that some basic parameters of model training such as definition of epochs number and model checkpoints are still accessible in ATI_Box. In addition, the ability to visualize model predictions through the quick image inspection feature provides insight into which structures the model has successfully learned and which remain a challenge. This allows one to better grasp the idea of semantic segmentation.

Beyond its analytical application, the platform can serve as a comprehensive educational environment for teaching convolutional neural networks implementation in image analysis. By integrating data upload, annotation, model training, inference, and quantitative evaluation within a single interface, it may enable students to observe the entire semantic segmentation pipeline in practice. Learners can create their own annotated datasets, configure class definitions, monitor training performance, and directly evaluate model using pixel-level IoU and object-wise metrics.

It has to be kept in mind that the usability of the presented version of the platform comes at a cost of its capability to adapt to specific problems. Therefore users have to be aware that some tasks may be solved more effectively with already existing dedicated tools. A good example here is a problem of segmentation of cells or of nuclei which are located in high proximity or even overlap. To solve this problem a very powerful tools exist of which StarDist [14] is worth mentioning. In its standard version StarDist is not a semantic segmentation tool i.e. it was not intended to multiclassification problems, but it is very efficient instance segmentation tool which allows separation of overlapping cells or nuclei.

The development process of ATI_Box supported by generative AI highlights a meta-level implication: generative AI may lower not only the barrier to applying deep learning methods, but also the barrier to constructing domain-specific AI infrastructure. ATI_Box therefore serves as a case study in AI-augmented scientific software engineering, illustrating how domain experts can create reproducible computational tools without direct institutional access to specialized engineering teams. This paradigm may have significant implications for the democratization of research-grade AI systems.

Author is aware of limitations of the platform. Although it was designed with usability in mind and aims to be accessible to non-expert users, its deployment still requires a certain level of technical proficiency due to its containerized (Docker-based) architecture. This limitation is at least partly mitigated by detailed tutorial ATI_Box installation provided in the supplementary material. The platform is designed for local deployment and its default configuration reflects this assumption. Service credentials for MinIO and Label Studio are hardcoded in the Docker Compose configuration to eliminate an additional setup step for the target user. Users who intend to host the platform on a network-accessible server must replace these default credentials with strong, unique values and restrict access to exposed service ports. Detailed instructions are provided in the repository documentation. Additionally, users should be aware that PyTorch model checkpoint files (.pth) can execute arbitrary code upon loading and should only be obtained from trusted sources.

The lack of functionality of 3D image analysis may be considered as important drawback by some users. As the platform was designed as a solution for entry-users it was thought that 2D analysis would be sufficient for this group. One of the functions that could prove useful but was not implemented is early stopping with model checkpointing which allows to stop training when model goes into overfitting and save the model with best convergence. This was intentionally omitted, for educational purposes. On one hand it could save time and resources but on the other it would obscure the idea of overfitting. To partially compensate for this potential disadvantage the training can be stopped at any time and model with weights effective from the last epoch saved.

## Supporting information

supplemenary file - installation guide

## Data availability

The datasets used in the Results section were acquired in the author’s laboratory and are available upon reasonable request. The platform can be tested with any user-provided images

## AI-use declaration

Claude (Anthropic, model version: claude-sonnet-4-6) was used to assist code generation. The final version was reviewed by the author.

## Code availability

The source code corresponding to this manuscript is available at: https://github.com/Przygodzkitom/semsegplat-local-full ATI_Box image is available at: https://github.com/Przygodzkitom?tab=packages

## Acknowledgements

The platform is released under the Apache 2.0 license, ensuring permissive reuse while remaining compatible with its core dependencies, including Label Studio and Streamlit (Apache 2.0), and MinIO (AGPL v3). MinIO runs as a separate container and ATI_Box’s code does not incorporate MinIO source code.

Author would like to thank to prof. Patrycja Przygodzka (Laboratory of Cellular and Molecular Biology, Institute of Medical Biology, Polish Academy of Sciences, 93-232 Lodz, Poland) and prof. Jacek Golański (Department of Haemostatic Disorders, Chair of Biomedical Sciences, Medical University of Lodz, ul. Mazowiecka 6/8, Lodz, 92-215, Poland) for providing images for scratch assay and blood smear test respectively.

