## Supplementary material for "ATI_Box: A Simple tool for convolutional neural network-based image semantic segmentation": supplemenary file - installation guide

#### ATI_Box Installation and First-Use Guide

This guide walks you through installing and running ATI_Box from scratch. No programming knowledge is required. The entire process takes approximately 20–40 minutes, most of which is waiting for a one-time download.

#### 1. What ATI_Box runs

ATI_Box is a set of three services that run together inside Docker, a programme that creates isolated lightweight virtual environments (called containers) on your computer. You interact with everything through your web browser. The three services are:

| Service | What it does | How you access it |
|---|---|---|
| Streamlit application | Main interface: upload images, train models, run inference, view results | http://localhost:8501 |
| Label Studio | Annotation tool: draw segmentation masks on your images | http://localhost:8080 |
| MinIO | File storage: keeps your images, annotations, and models organised | http://localhost:9001 (admin console) |

All your data (images, annotations, trained models) is stored in ordinary folders on your computer, not locked inside the containers. This means your work is safe even if you update or reinstall the platform.

#### 2. System requirements

| Component | Requirement |
|---|---|
| RAM | 8 GB minimum, 16 GB recommended for training |
| Disk space | 15 GB free (Docker images + your data) |
| Internet | Required for initial download only (~5–7 GB, one time) |
| GPU | Optional. NVIDIA GPU with CUDA speeds up training significantly, but CPU works |
| Operating system | Windows 10/11, macOS, or Linux |

#### 3. Install Docker

Docker is the only software you need to install. It creates the environment in which ATI_Box runs.

##### 3.1. Windows

Download and install **Docker Desktop for Windows** from https://docs.docker.com/desktop/install/windows-install/. During installation, Docker will ask you to enable WSL 2 (Windows Subsystem for Linux) — accept this, it is required. After installation, open Docker Desktop from the Start menu. Wait until the whale icon in the taskbar stops animating (this means Docker is ready).

> *Tip: If Docker Desktop asks you to restart your computer during installation, do so before continuing. If Docker Desktop fails to start or displays an error mentioning “hardware virtualisation” or “Hyper-V”, you need to enable CPU virtualisation in your computer’s BIOS/UEFI settings. This feature is called Intel VT-x (on Intel processors) or AMD-V / SVM (on AMD processors). To access the BIOS, restart your computer and press the key shown briefly on the boot screen (typically F2, F10, F12, Del, or Esc — the exact key depends on your computer manufacturer). Inside the BIOS, look under a menu called “Advanced”, “CPU Configuration”, or “Security” for a setting named “Virtualisation Technology”, “VT-x”, or “SVM Mode”, set it to Enabled, save, and restart. On most modern computers this setting is already enabled, but some manufacturers ship it turned off by default.*

##### 3.2. macOS

Download and install **Docker Desktop for Mac** from https://docs.docker.com/desktop/install/mac-install/. After installation, open Docker Desktop. Wait for the whale icon in the menu bar to stop animating.

##### 3.3. Linux

Install Docker Engine and the Compose plugin following the official instructions for your distribution at https://docs.docker.com/engine/install/. After installation, you must add your user to the docker group so you can run Docker commands without typing sudo every time:

```
sudo usermod -aG docker $USER
```

Then log out of your computer session and log back in. Verify the installation by opening a terminal and typing:

```
docker --version
docker compose version
```

Both commands should print version numbers without errors.

#### 4. GPU support (optional)

**Skip this section entirely** if you do not have an NVIDIA GPU or are unsure. ATI_Box works on CPU — training simply takes longer (hours instead of minutes). You can always add GPU support later.

If you have an NVIDIA GPU and want to use it for faster training, you need to install the NVIDIA Container Toolkit so that Docker can access your GPU. On Linux, run the following commands in a terminal:

```
curl -fsSL https://nvidia.github.io/libnvidia-container/gpgkey \
  | sudo gpg --dearmor -o /usr/share/keyrings/nvidia-container-toolkit-keyring.gpg


curl -s -L https://nvidia.github.io/libnvidia-container/stable/deb/nvidia-container-toolkit.list \
  | sed 's#deb https://#deb [signed-by=/usr/share/keyrings/nvidia-container-toolkit-keyring.gpg] https://#g' \
  | sudo tee /etc/apt/sources.list.d/nvidia-container-toolkit.list
```

```
sudo apt-get update
sudo apt-get install -y nvidia-container-toolkit
sudo nvidia-ctk runtime configure --runtime=docker
sudo systemctl restart docker
```

On Windows, GPU access works automatically through WSL 2 as long as you have the latest NVIDIA driver installed. On macOS, GPU acceleration is not available; training runs on CPU.

### 5. Download the startup files

You need to download a small set of configuration files that tell Docker how to run ATI_Box. These are not the application itself (that is downloaded automatically by Docker in the next step), just the instructions for it.

#### 5.1. Option A: Download from the terminal

First, create a folder for your project and navigate into it. Once in your folder Open a terminal (or PowerShell on Windows) and type the following commands.

**Linux / macOS:**

```
curl -O https://raw.githubusercontent.com/Przygodzkitom/semsegplat-local-full/main/docker-compose.yml
curl -O https://raw.githubusercontent.com/Przygodzkitom/semsegplat-local-full/main/docker-compose.gpu.yml
curl -O https://raw.githubusercontent.com/Przygodzkitom/semsegplat-local-full/main/start.sh
chmod +x start.sh
```

**Windows (PowerShell):**

```
Invoke-WebRequest -Uri "https://raw.githubusercontent.com/Przygodzkitom/
  semsegplat-local-full/main/docker-compose.yml" -OutFile docker-compose.yml
Invoke-WebRequest -Uri "https://raw.githubusercontent.com/Przygodzkitom/
  semsegplat-local-full/main/docker-compose.gpu.yml" -OutFile docker-compose.gpu.yml
Invoke-WebRequest -Uri "https://raw.githubusercontent.com/Przygodzkitom/
  semsegplat-local-full/main/start.bat" -OutFile start.bat
```

#### 5.2. Option B: Download manually from GitHub

If you prefer not to use the terminal for this step, visit the repository at https://github.com/Przygodzkitom/semsegplat-local-full and download the following files: docker-compose.yml, docker-compose.gpu.yml, and start.sh (Linux/macOS) or start.bat (Windows). Place all downloaded files in a single folder (for example, a folder called semseg-platform on your Desktop).

### 6. Start the platform

**Before running:** make sure Docker is running. On Windows and macOS, open Docker Desktop and wait for the whale icon to stop animating. On Linux, you can check by typing `docker ps` in a terminal — if it prints a table (even an empty one) without an error, Docker is running.

Open a terminal, navigate to the folder where you placed the startup files, and run:

**Linux / macOS:**

```
./start.sh
```

**Windows (PowerShell):**

```
.\start.bat
```

> *Important: In PowerShell, you must type .\start.bat with the dot and backslash. Typing just start.bat runs a different Windows command.*

On the first run, Docker will download the application images from the registry. This is a one-time download of approximately 5–7 GB and may take 10–20 minutes depending on your internet connection. Subsequent starts take only a few seconds.

The startup script automatically creates the necessary data folders on your computer, detects whether you have a GPU, and launches the platform with the correct configuration. When you see messages indicating that Streamlit and Label Studio have started, the platform is ready.

#### 7. Open the platform in your browser

Once the startup messages indicate the services are running, open your web browser and go to:

| Application | Address | Login credentials |
|---|---|---|
| Streamlit (main interface) | http://localhost:8501 | None required |
| Label Studio (annotation) | http://localhost:8080 |<br>Password: admin |
| MinIO console (storage) | http://localhost:9001 | Username: minioadmin<br>Password: minioadmin123 |

> *Note: On the very first start, Label Studio takes an extra 30–60 seconds to initialise its database. If you see a loading screen at http://localhost:8080, wait a moment and refresh the page.*

#### 8. Your first workflow: from images to results

This section walks through the entire pipeline to help you verify that everything is working and to familiarise you with the interface.

##### 8.1. Upload images

Open http://localhost:8501 in your browser. In the Streamlit interface, navigate to the Upload section. Select and upload the images you want to analyse. The images are automatically stored in MinIO. You can upload images in TIFF, JPEG, or PNG format; non-PNG images are converted automatically.

##### 8.2. Annotate

In the Streamlit interface, define the number and names of your segmentation classes (for example: "normal cells" and "apoptotic cells") and choose an annotation mode (polygon or brush). Then click the button to open Label Studio. In Label Studio, log in with the default credentials ( / admin), open the project that was created automatically, and draw segmentation masks on your images. When you are finished annotating, return to the Streamlit interface.

##### 8.3. Train a model

In the Streamlit interface, go to the Training section. Choose the number of training epochs (the default of 100 is a reasonable starting point). Click the button to start training. The interface will show you the progress and display training and validation loss curves as the model learns. Training time depends on the number of images, their size, and whether you are using a GPU. As a rough guide: with a GPU, expect 15–30 minutes; on CPU, expect 2–4 hours for a typical dataset of 20–30 images.

##### 8.4. Evaluate the model

After training completes, you can evaluate the model in two ways. First, use the single-image inference mode to upload a new image and visually inspect the predicted segmentation masks. Adjust the probability threshold to see how it affects the results. Second, use the ground-truth evaluation mode, which automatically runs the model on the held-out test images (set aside before training) and reports pixel-wise IoU and object-wise precision, recall, and F1-score.

##### 8.5. Batch analysis

When you are satisfied with the model, go to the Batch Analysis section. Upload a set of images from your experiment. The platform processes them automatically and produces a table with the number of detected objects and the area covered by each class for every image. You can export the results as a CSV file for further analysis in statistical software.

#### 9. Stopping and restarting

To stop the platform, open a terminal, navigate to your semseg-platform folder, and type:

```
docker compose down
```

Your data (images, annotations, trained models) is preserved on your computer. To start the platform again later, simply run the start script again:

```
./start.sh          # Linux / macOS
.\start.bat         # Windows (PowerShell)
```

#### 10. Updating to a newer version

When a new version of ATI_Box is released, you can update by pulling the latest images. Open a terminal in your semseg-platform folder and run:

```
docker compose pull
docker compose up -d
```

Your data folders (minio-data, label-studio-data, models) are not affected by the update.

#### 11. Troubleshooting

If something does not work as expected, the following table lists the most common issues and their solutions.

| Problem | Likely cause | Solution |
| --- | --- | --- |
| "Permission denied" on /var/run/docker.sock (Linux) | Your user is not in the docker group | Run: sudo usermod -aG docker $USER — then log out and back in |
| "Port already in use" error | Another programme is using port 8501, 8080, 9000, or 9001 | Stop the other programme, or edit docker-compose.yml and change the port number on the left side of the colon |
| Docker commands fail (macOS / Windows) | Docker Desktop is not running | Open Docker Desktop and wait for it to finish starting |
| Docker Desktop error mentioning “hardware virtualisation” or “Hyper-V” | CPU virtualisation is disabled in BIOS | Restart computer, enter BIOS (F2/F10/Del/Esc at boot), find Virtualisation Technology / VT-x / SVM Mode under Advanced or CPU Configuration, set to Enabled, save and restart |
| GPU not detected inside the container | NVIDIA Container Toolkit not installed or Docker not restarted | Revisit Section 4 above; after installing the toolkit run: sudo systemctl restart docker |
| Label Studio shows blank page | Database is still initialising | Wait 60 seconds and refresh the page |
| MinIO connection error in Streamlit | MinIO has not finished starting | Wait 30 seconds after starting the platform and refresh |
| Out of memory during training | Insufficient RAM or VRAM | Close other applications; on Docker Desktop increase memory in Settings → Resources |

#### 12. Backing up your data

All your work is stored in three folders inside the semseg-platform directory: minio-data (your images and annotations), label-studio-data (annotation project database), and models/checkpoints (trained model files). To create a backup, simply copy these three folders to a safe location. To restore, copy them back into place before starting the platform.

> *Important: Always stop the platform (docker compose down) before backing up or restoring data. Copying files while the services are running can corrupt the database.*
